# Disruption of the VEGFR3–HSPG2 induces lymphatic reflux dysfunction and sepsis

**DOI:** 10.64898/2026.01.28.702461

**Authors:** Yiming Zhang, Linfeng Liu, Wei Guo, Shunshun Wang, Xinyi Wu, Yanxu Jiang, Pu-hong Zhang, Jianguang Wang, Shengwei Jin

**Affiliations:** Wenzhou Medical University; the Second Affiliated Hospital and Yuying Children's Hospital of Wenzhou Medical University; Wenzhou Medical University, Wenzhou, Zhejiang, China; Second Affiliated Hospital of Wenzhou Medical University

## Abstract

The lymphatic system plays a central role in fluid transport and immune regulation, yet its contribution to sepsis remains incompletely understood. Analysis of more than 500,000 individuals from the UK Biobank reveals that lymphatic system disorders and genetic variants in FLT4 (VEGFR3) are strongly associated with increased sepsis risk. Single-cell RNA sequencing further shows a marked reduction of VEGFR3^high lymphatic endothelial cells during sepsis. LEC-specific deletion of VEGFR3 impairs lymphatic reflux without disrupting vessel architecture, leading to barrier failure, systemic inflammation, bacterial dissemination, and sepsis. Mechanistically, VEGFR3 sustains lymphatic glycocalyx integrity by promoting SOX18-dependent HSPG2 transcription and heparan sulfate biosynthesis. Consistently, LEC-specific HSPG2 knockdown or pharmacological inhibition of SOX18 phenocopies VEGFR3 deficiency, whereas heparan sulfate supplementation alleviates sepsis-like pathology. Moreover, HSPG2 genetic variants are significantly associated with both lymphatic disorders and sepsis. Collectively, these findings identify lymphatic reflux dysfunction as a pathogenic determinant of sepsis and establish disruption of the VEGFR3–SOX18–HSPG2–heparan sulfate axis as an active driver of disease.

## Introduction

The lymphatic system is a fundamental vascular network that maintains tissue fluid homeostasis, immune surveillance and regulation^[1]^. Lymphatic endothelial cells(LECs) form the structural and functional foundation of lymphatic vessels and are essential for barrier integrity, fluid drainage^[2, 3]^. When lymphatic vessels function is compromised, impaired clearance of interstitial fluid and inflammatory mediators leads to tissue edema and amplification of inflammatory responses^[4]^. Such tissue edema has been increasingly implicated in the pathogenesis of inflammatory disorders and may contribute to the progression toward systemic inflammation, highlighting the essential role of lymphatic vessels function^[5, 6]^.

Vascular endothelial growth factor receptor 3(VEGFR3, also known as FLT4) is a central signaling receptor that governs lymphatic endothelial identity and function^[7]^. Upon activation by its ligand VEGF-C, VEGFR3 initiates downstream signaling pathways, including PI3K – AKT and MAPK cascades, that regulate lymphatic endothelial cell survival, migration, and barrier stability^[8–11]^. Genetic studies have established that VEGFR3 is indispensable for lymphatic development, as its global loss results in failed lymphangiogenesis and embryonic lethality, while perturbation of VEGFR3 signaling disrupts lymphatic endothelial growth and function^[12, 13]^. Notably, although VEGFR3 has been extensively studied for its essential role in lymphatic endothelial cell proliferation and lymphangiogenesis, far less is known about its involvement in non-proliferative aspects of lymphatic endothelial function, particularly in the regulation of the endothelial glycocalyx and associated polysaccharide structures.

Effective lymphatic reflux depends not only on lymphatic vessel architecture but also on the luminal microenvironment that supports fluid transport and endothelial stability. Multiple factors can influence lymphatic reflux efficiency, including endothelial barrier integrity, mechanical sensing, and the molecular composition of the lymphatic lumen. A prominent structural feature of lymphatic vessels is the glycocalyx, a polysaccharide-rich coating lining the endothelial surface that modulates permeability, mechanotransduction, and inflammatory signaling^[14–16]^. Disruption of this luminal glycocalyx compromises lymphatic fluid transport and promotes tissue edema and inflammatory amplification^[17, 18]^. Among glycocalyx-associated polysaccharides, heparan sulfate(HS) plays a particularly important role due to its high negative charge density and structural versatility, which support glycocalyx integrity and fluid homeostasis within lymphatic vessels^[19, 20]^. HS biosynthesis and organization critically depend on specific core proteins, most notably heparan sulfate proteoglycan 2(HSPG2, perlecan), which provides the scaffold required for HS assembly and function^[21–24]^. Despite the critical role of HS in maintaining lymphatic transport capacity, the regulatory mechanisms governing HS biosynthesis in lymphatic endothelial cells, and their contribution to lymphatic reflux and inflammatory pathology still require further investigation.

In this study, we identify a VEGFR3 – SOX18 – HSPG2 signaling axis that governs lymphatic glycocalyx integrity and lymphatic reflux, and demonstrate that disruption of this pathway induces tissue injury and sepsis. These findings establish lymphatic dysfunction as an active contributor to sepsis pathogenesis and reveal a mechanistic framework linking lymphatic endothelial regulation to sepsis.

## Results

### 1. Lymphatic dysfunction is a major risk factor for sepsis and is closely associated with VEGFR3 function

To investigate whether lymphatic dysfunction contributes to the pathogenesis of sepsis, we first analyzed data from 502,137 participants in the UK Biobank(Figure 1A) and adjusted for age, sex, and other covariates(Supplementary Fig.1A). Logistic regression revealed that individuals with lymphatic system disorders had a 5.4136-fold higher risk of developing sepsis, compared with controls(Figure 1B; Supplementary Fig.1B). Notably, this risk was even greater among individuals with hereditary lymphoedema(Figure 1C; Supplementary Fig.1C), indicating a strong association between lymphatic impairment and sepsis.

**Figure 1.**
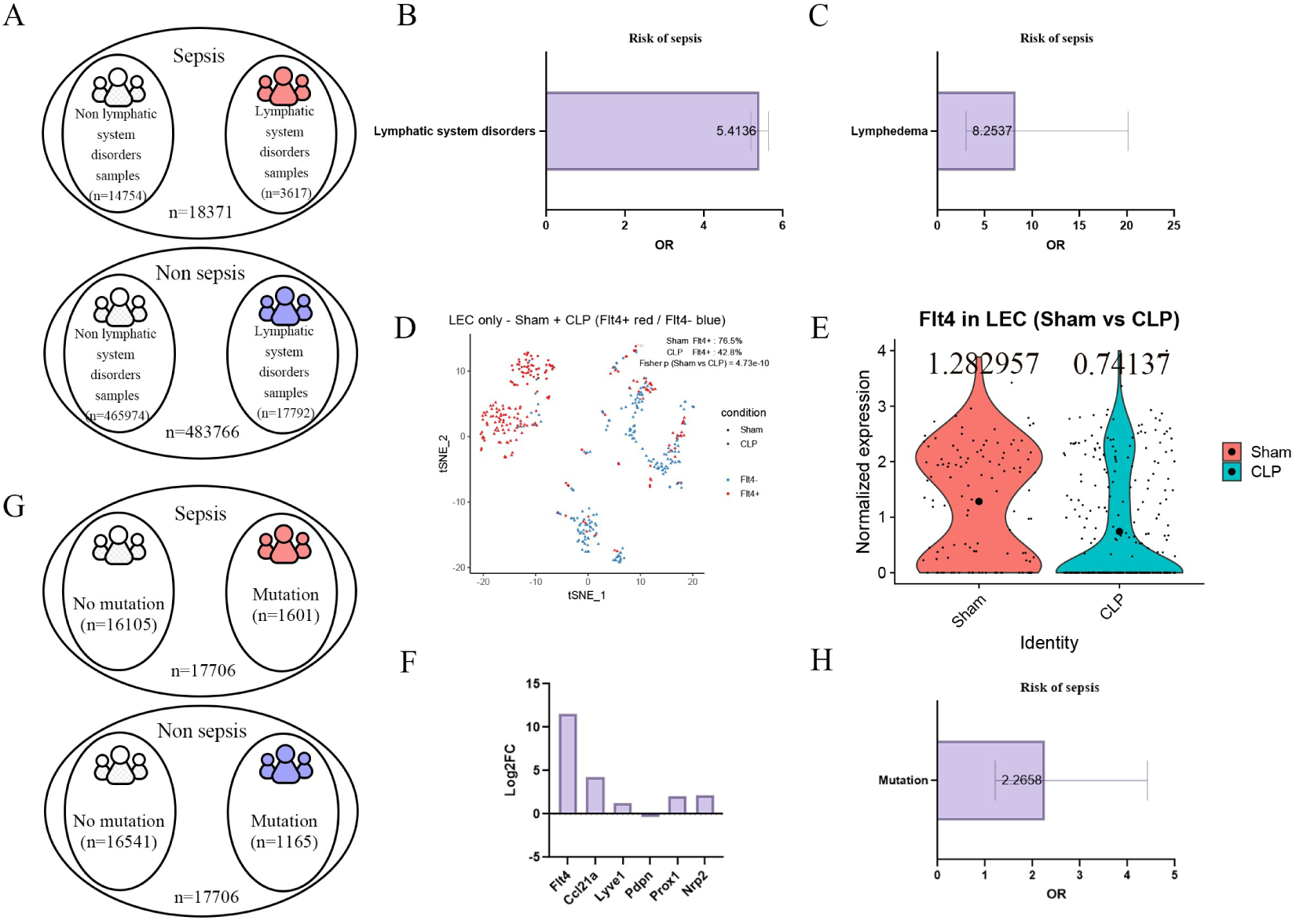
Lymphatic system disorders and VEGFR3 genetic alterations are associated with sepsis in human cohorts and mouse single-cell datasets. **(A)** Schematic overview of the UK Biobank cohort(n = 502,137). White indicates the control population, red and blue indicates patients diagnosed with lymphatic system disorders in sepsis and non sepsis group. **(B)** Logistic regression showing a significantly increased sepsis incidence in individuals with lymphatic system disorders(OR = 5.41). **(C)** Primary lymphedema patients exhibit an even higher sepsis risk(OR = 8.25). **(D)** UMAP plot of scRNA-seq data demonstrating reduced proportions of VEGFR3⁺ LECs in CLP-induced septic mice. Red denotes FLT4^high LECs and blue denotes FLT4^low LECs; circles represent sham-operated mice and triangles represent CLP-induced mice. **(E)** FLT4 expression is significantly reduced in LECs following CLP-induced sepsis. **(F)** Among multiple LEC markers, FLT4 shows the most pronounced decrease in expression. **(G)** Population-level analysis showing higher frequency of FLT4 variants among sepsis patients compared with controls. White indicates individuals without FLT4 mutations, red indicates sepsis patients carrying FLT4 mutations, and blue indicates non-sepsis individuals carrying FLT4 mutations. (H) Individuals carrying FLT4 variants show a significantly increased risk of sepsis(OR = 1.4093). Data are presented as odds ratios(OR) with 95% confidence intervals(CIs).

To further explore the molecular basis of this association, we analyzed the CLP sepsis single-cell dataset GSE207651. In septic mice, the proportion of VEGFR3-high LECs was markedly reduced(Figure 1D), and their mean expression level was significantly decreased(Figure 1E–F). Consistent with these findings, among 35,414 individuals with available genomic data, the proportion of VEGFR3 variant carriers was significantly higher in sepsis cases(9.04%) compared with non-sepsis controls(6.58%)(Figure 1G–H; Supplementary Fig.1D–E). Together, these population-based and single-cell analyses strongly suggest that VEGFR3 dysfunction affects lymphatic homeostasis and significantly increases the risk of sepsis.

### 2. LEC-specific deletion of VEGFR3 induces lymphatic reflux dysfunction and sepsis

To investigate the functional role of VEGFR3 in LECs, we generated Prox1CreERT2-mediated LEC-specific VEGFR3 knockout mice(Figure 2A; Supplementary Figure 2). After intraperitoneal administration of tamoxifen, VEGFR3^iΔ LEC mice exhibited significantly reduced survival, but the lymphatic vessel structure remained intact(Figure 2B, Supplementary Figure 2B). These mice displayed markedly elevated circulating levels of pro-inflammatory cytokines, including TNF-α and IL-6(Figure 2C), together with increased systemic bacterial burden(Figure 2D).

**Figure 2.**
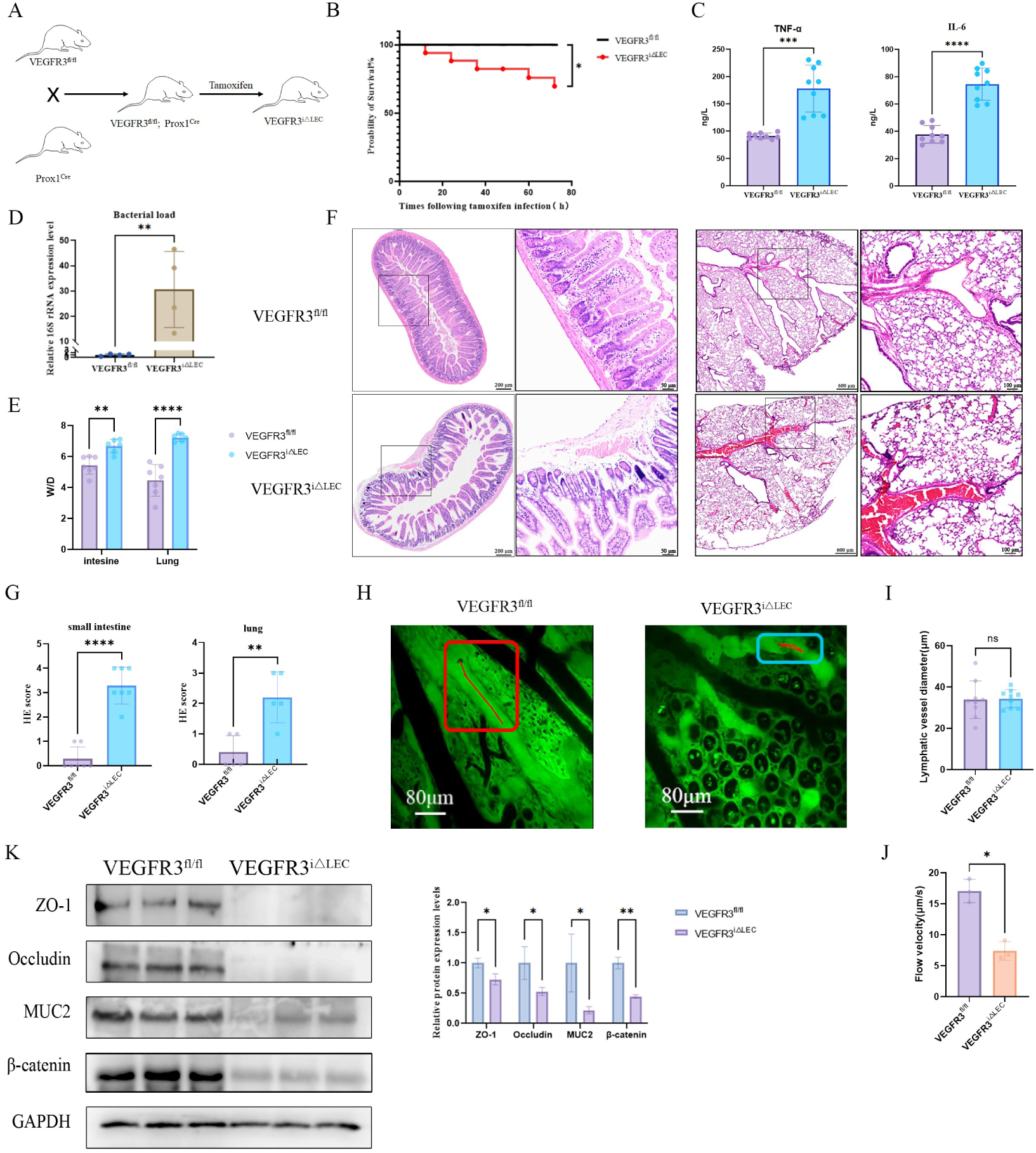
LEC-specific VEGFR3 deletion impairs lymphatic reflux and induces sepsis. **(A)** Generation strategy of Prox1CreERT2;VEGFR3^fl/fl(VEGFR3^iΔLEC) mice. **(B)**Kaplan–Meier survival analysis showing reduced survival following tamoxifen-induced VEGFR3 deletion(n=18).**(C–D)** Significantly increased plasma TNF-α/IL-6(n=9) and elevated bacterial load(n=4). **(E)** Increased tissue water content(W/D ratio) in intestine and lung(n=7). **(F)** Representative H&E staining showing disrupted intestinal mucosa and alveolar damage. **(G)** Histological injury scoring for gut and lung. **(H)** In vivo lymphatic imaging showing impaired lymphatic reflux in VEGFR3^iΔLEC mice. The curve marked in the box represents the movement trajectory of cells in the lymphatic vessels. **(I)** Lymphatic vessel diameter. **(J)** Cell flow rate within lymphatic vessels. **(K)** Relative expression of intestinal barrier proteins(n=3). Data are presented as mean ± SEM. *P < 0.05, **P < 0.01, ***P < 0.001, ****P < 0.0001; ns, not significant.

Both intestinal and pulmonary tissues from VEGFR3^iΔLEC mice exhibited significantly increased water content(W/D) (Figure 2E). Histological analysis revealed marked structural disruption, including loss of goblet cells, and disorganized smooth muscle layers in the intestine, together with alveolar rupture, fusion, and inflammatory infiltration in the lungs(Figure 2F). Injury scores confirmed more severe intestinal and pulmonary damage(Figure 2G), indicating exacerbated tissue edema. In vivo lymphatic imaging further demonstrated marked impairment of lymphatic reflux in VEGFR3^iΔLEC mice(Figure 2H; Video 1). After statistical analysis, we found that the diameter of the lymphatic vessels did not change significantly(Figure 2I), lymph flow velocity was markedly reduced(Figure 2J). Consistent with these functional defects, expression levels of epithelial and junctional proteins, including ZO-1, occludin, MUC2, and β-catenin, were substantially decreased(Figure 2K), indicating compromised barrier integrity. Collectively, these findings indicate that VEGFR3 is essential for maintaining lymphatic reflux and epithelial barrier integrity, and that its disruption predisposes to tissue injury and sepsis.

### 3. VEGFR3 knockdown disrupts the HSPG2-dependent heparan sulfate synthesis system in LECs

To define the molecular mechanisms by which VEGFR3 regulates lymphatic endothelial function, we performed transcriptomic profiling following VEGFR3 knockdown in human lymphatic endothelial cells(HLECs). Differential expression analysis revealed extensive transcriptional reprogramming upon VEGFR3 silencing(Figure 3A-B). Pathway enrichment analysis showed that HS biosynthesis pathways were significantly downregulated following VEGFR3 silencing(Figure 3C–D). Gene set enrichment analysis(GSEA) further confirmed global suppression of HS–GAG metabolism(Figure 3E), suggesting a potential link between VEGFR3 signaling and glycocalyx integrity.

**Figure 3.**
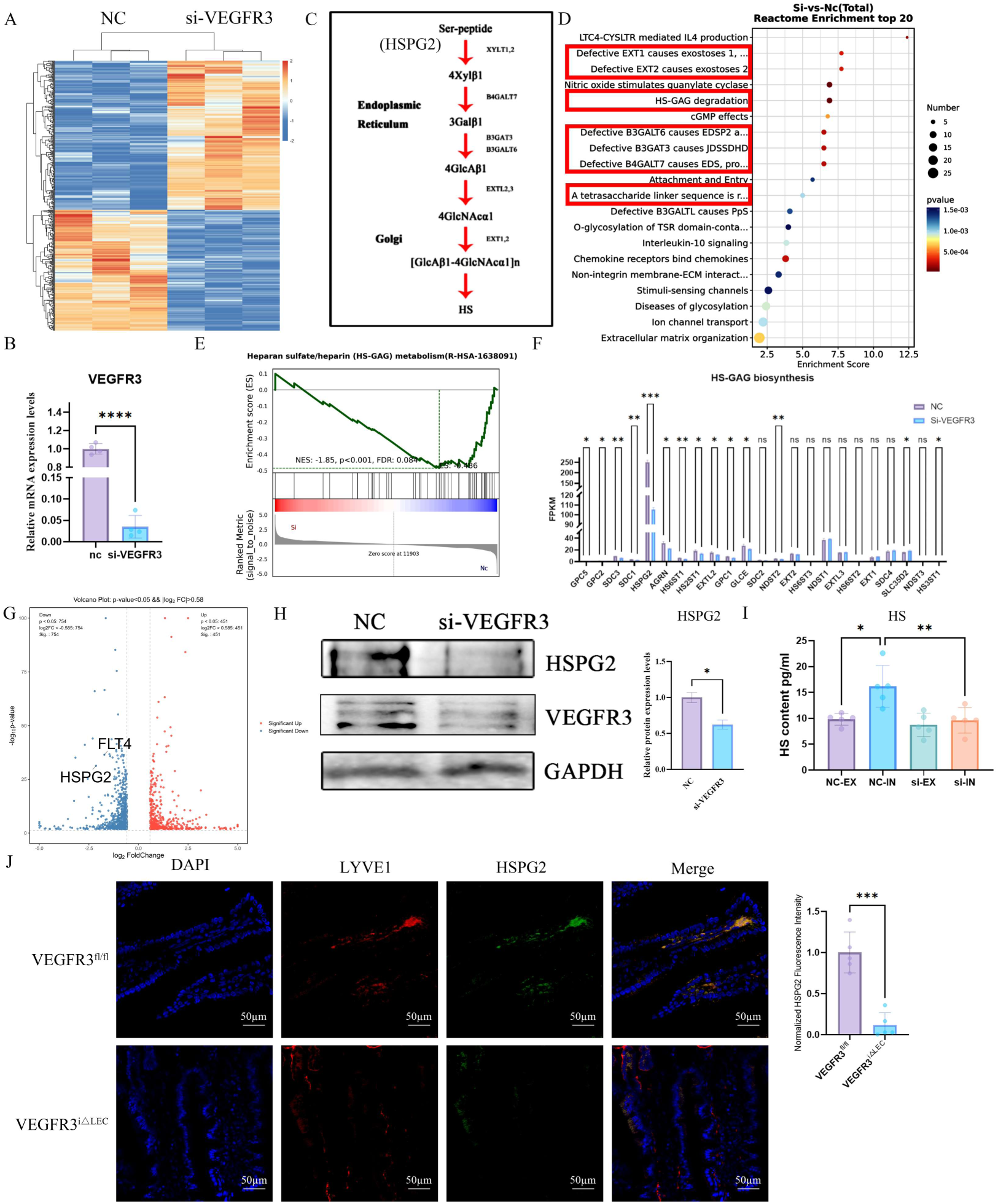
VEGFR3 loss disrupts HSPG2-dependent heparan sulfate biosynthesis. **(A)** Transcriptomic profiling of VEGFR3-deficient HLECs. **(B)** Validation of omics results. **(C)** Synthetic pathways of HS. **(D)** Reactome pathway enrichment analysis following VEGFR3 knockdown. The red boxes mark the pathways related to HS, most of which correspond to figure 3C. **(E)** GSEA confirms suppression of HS–GAG pathway. **(F)** Differential expression analysis identifying HSPG2 as the most significantly reduced HS-core gene. **(G)** Volcano plot. **(H)**Relative expression of HSPG2(n=3). **(I)** ELISA detection showed that knocking down VEGFR3 resulted in a decrease in intracellular HS, while the content of free HS remained unchanged(n=5). NC-EX: negative control-extracellular; NC-IN: negative control-intracellular; si-EX: siVEGFR3-extracellular; si-IN: siVEGFR3-intracellular. **(J)** Immunofluorescence staining confirming loss of HSPG2 expression in lymphatic vessels of intestinal villi.Data are presented as mean ± SEM. *P < 0.05, **P < 0.01, ***P < 0.001, ****P < 0.0001; ns, not significant.

Among all HS-related genes, HSPG2 exhibited the most significant VEGFR3-dependent reduction(Figure 3F–G). At the protein level, immunoblotting confirmed a substantial decrease in HSPG2 protein abundance in si-VEGFR3 cells(Figure 3H). Consistent with these findings, intracellular HS levels were markedly reduced in si-VEGFR3 cells, whereas extracellular HS showed no significant change(Figure 3I), indicating a specific impairment in HS biosynthesis rather than global secretion defects. Immunofluorescence staining further showed that the expression level of HSPG2 on the lymphatic vessels in the intestinal villi was significantly reduced(Figure 3J), confirming that VEGFR3-dependent regulation of HSPG2 occurs in vivo. Together, these findings identify VEGFR3 as a central upstream regulator of the HSPG2 – HS axis and establish its essential role in maintaining lymphatic glycocalyx integrity and endothelial homeostasis.

### 4. LEC-specific HSPG2 deficiency impairs lymphatic reflux and induces sepsis

To determine whether HSPG2 functions as a critical downstream effector of VEGFR3 signaling, we generated LEC-specific HSPG2 knockdown mice using AAV-shHSPG2 in Prox1CreERT2 mice(Figure 4A; Supplementary Fig.3). HSPG2 deficiency significantly elevated plasma TNF-α and IL-6(Figure 4B), accompanied by a marked increase in systemic bacterial burden(Figure 4C), markedly elevated intestinal and pulmonary W/D ratios(Figure 4D). Lymphatic imaging revealed profound impairment of lymphatic flow without significant changes in vessel diameter(Figure 4E–G; Video 2). Histopathological analysis demonstrated extensive epithelial injury, including villus blunting and epithelial disruption in the intestine, as well as diffuse inflammatory infiltration and alveolar damage in the lung(Figure 4H-I). Collectively, these findings indicate that HSPG2 is a critical downstream effector of VEGFR3 signaling required for maintaining lymphatic transport and barrier integrity. Loss of HSPG2 recapitulates key pathological features observed upon VEGFR3 deficiency, leading to lymphatic dysfunction, inflammatory amplification, and systemic tissue injury.

**Figure 4.**
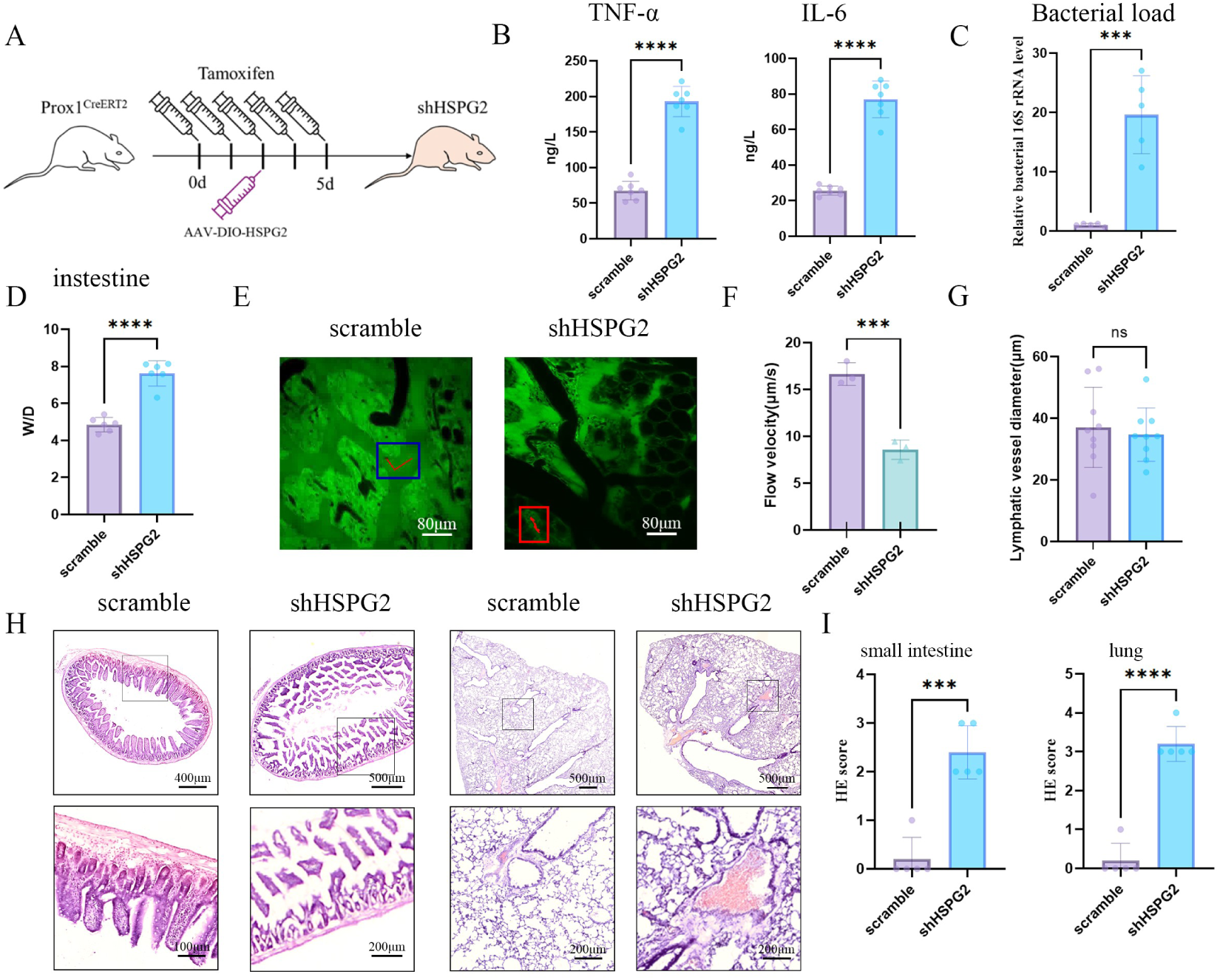
LEC-specific HSPG2 knockdown impairs lymphatic reflux and induces sepsis. **(A)** AAV-shHSPG2 delivery strategy in Prox1CreERT2 mice. **(B-C)** Increased TNF-α/IL-6(n=7) and bacterial burden(n=5) after HSPG2 silencing. **(D)** Elevated W/D ratio indicating aggravated edema(n=6). **(E–F)** In vivo lymphatic imaging revealing impaired lymphatic drainage without significant lymphatic vessel dilation. The curve marked in the box represents the movement trajectory of cells in the lymphatic vessels per unit time. **(G)** The diameter of the lymphatic vessels. **(H)** H&E staining. **(I)** injury scoring of intestinal and lung. Data are presented as mean ± SEM. *P < 0.05, **P < 0.01, ***P < 0.001, ****P < 0.0001; ns, not significant.

### 5. VEGFR3 regulates HSPG2 transcription through SOX18 activation

To elucidate the molecular mechanism by which VEGFR3 regulates HSPG2 expression, we first performed integrative transcriptional analyses following VEGFR3 knockdown. A Sankey diagram was generated to visualize transcription factors whose expression was altered upon VEGFR3 knockdown(Figure 5A). Using integrative bioinformatic analyses, we evaluated whether these transcription factors are expressed in LECs, whether they possess predicted binding sites within the HSPG2 locus, whether they associate with sepsis, and whether they respond to VEGFR3 signaling(Figure 5B, Supplementary figure 4). Among these candidates, SOX18 emerged as the most strongly correlated transcription factor, displaying both reduced expression under VEGFR3 deficiency and a strong association with HSPG2 expression.

**Figure 5.**
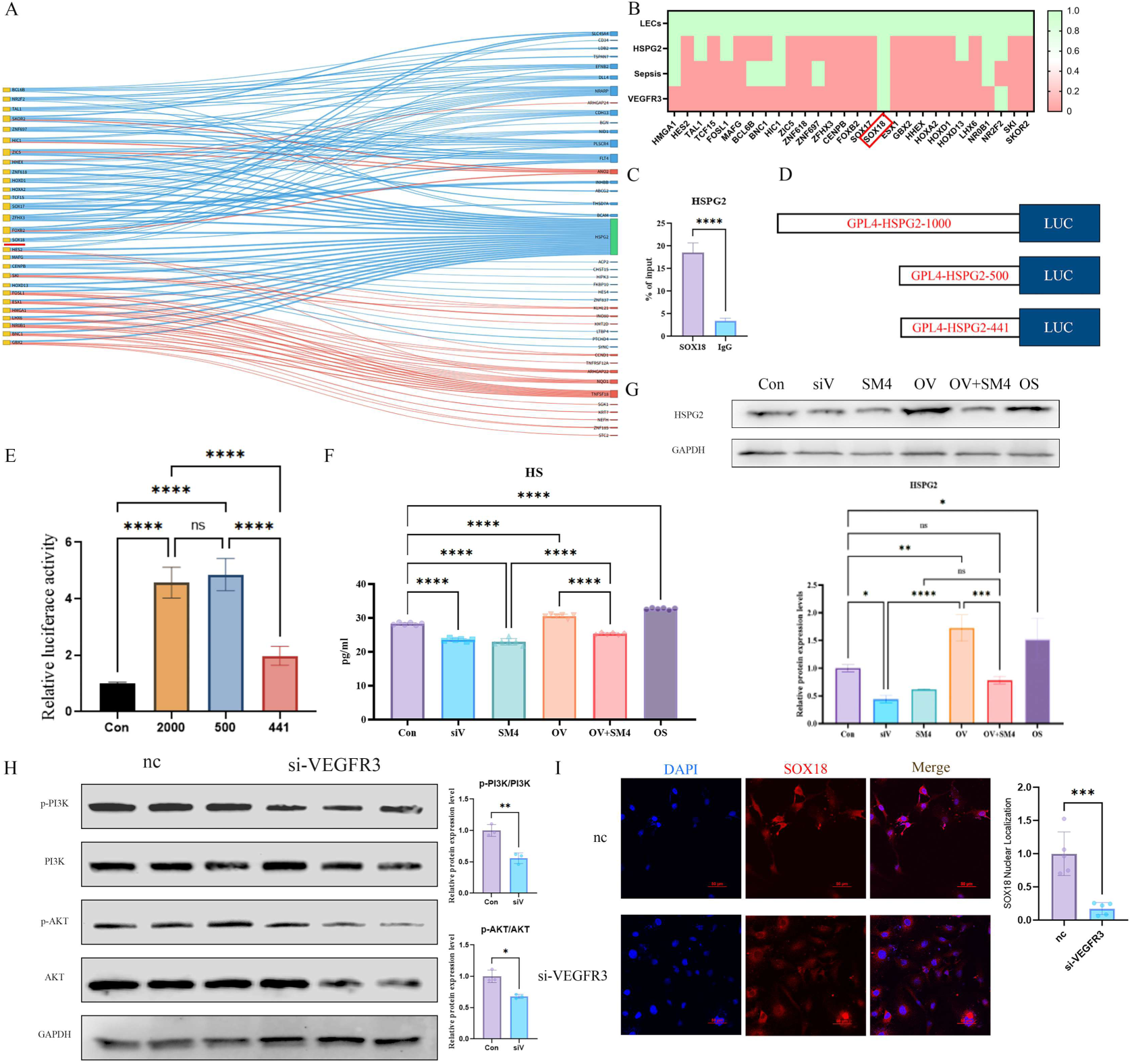
VEGFR3 regulates HSPG2 transcription through PI3K–AKT–dependent activation of SOX18. **(A)** The Sankey diagram illustrates the changes in transcription factors affected by VEGFR3 knockdown, which can be used to screen for key factors that may regulate HSPG2. **(B)** Bioinformatics integrated analysis was conducted on the expression of these transcription factors in LECs, their potential binding to HSPG2, their association with sepsis, and whether they are regulated by VEGFR3. **(C)** ChIP-qPCR demonstrating SOX18 binding to the HSPG2 promoter. **(D-E)** Luciferase reporter constructs locate the core SOX18 binding region at −441/−500 bp. **(F)** HS level detection(n=6). VEGFR3 knockdown(siV); SOX18 inhibitor(SM4); SOX18 overexpression(OS); VEGFR3 overexpression(OV). **(G)** Western blot(n=3). **(H)** VEGFR3 deficiency inhibited PI3K and AKT phosphorylation(n=3). **(I)** Confocal imaging showing impaired nuclear localization of SOX18 in VEGFR3-deficient HLECs(n=5). Data are presented as mean ± SEM. *P < 0.05, **P < 0.01, ***P < 0.001, ****P < 0.0001; ns, not significant.

To determine whether SOX18 directly regulates HSPG2 transcription, we performed assays, which demonstrated significant enrichment of SOX18 at the HSPG2 promoter region(Figure 5C). To identify the functional regulatory region, we constructed luciferase reporters containing 1000 bp, 500 bp, and 441 bp fragments of the HSPG2 promoter(Figure 5D). Luciferase assays revealed that the 441–500 bp region exhibited the highest transcriptional activity and was highly sensitive to SOX18 regulation(Figure 5E), indicating that this interval contains the core SOX18 binding motif.

Consistent with these findings, silencing of VEGFR3 or pharmacological inhibition of SOX18 markedly reduced HSPG2 and HS expression, whereas enforced expression of VEGFR3 or SOX18 significantly increased HSPG2 and HS levels(Figure 5F-G). Notably, overexpression of VEGFR3 failed to restore HSPG2 expression when SOX18 transcriptional activity was inhibited, indicating that SOX18 functions downstream of VEGFR3 in regulating HSPG2 transcription.

At the signaling level, VEGFR3 knockdown resulted in marked suppression of PI3K and AKT phosphorylation(Figure 5H). Immunofluorescence analysis further demonstrated that VEGFR3 knockdown significantly reduced SOX18 nuclear localization(Figure 5I), while the PI3K/AKT signaling pathway plays a key role in transcription factor nuclear translocation. Collectively, these data demonstrate that VEGFR3 promotes SOX18 nuclear entry via PI3K–AKT signaling, thereby driving HSPG2 transcription and maintaining HS biosynthesis and lymphatic endothelial homeostasis.

### 6. The VEGFR3/SOX18/HSPG2 axis regulates lymphatic reflux and sepsis

To further examine the functional role of the VEGFR3–SOX18–HSPG2 axis in vivo, mice were treated with the SOX18 inhibitor SM4. SM4 administration elicited a pronounced systemic inflammatory response, as evidenced by significantly elevated plasma levels of TNF-α and IL-6 compared with control mice(Figure 6A). Concomitantly, SM4-treated mice exhibited a marked increase in bacterial burden(Figure 6B).

**Figure 6.**
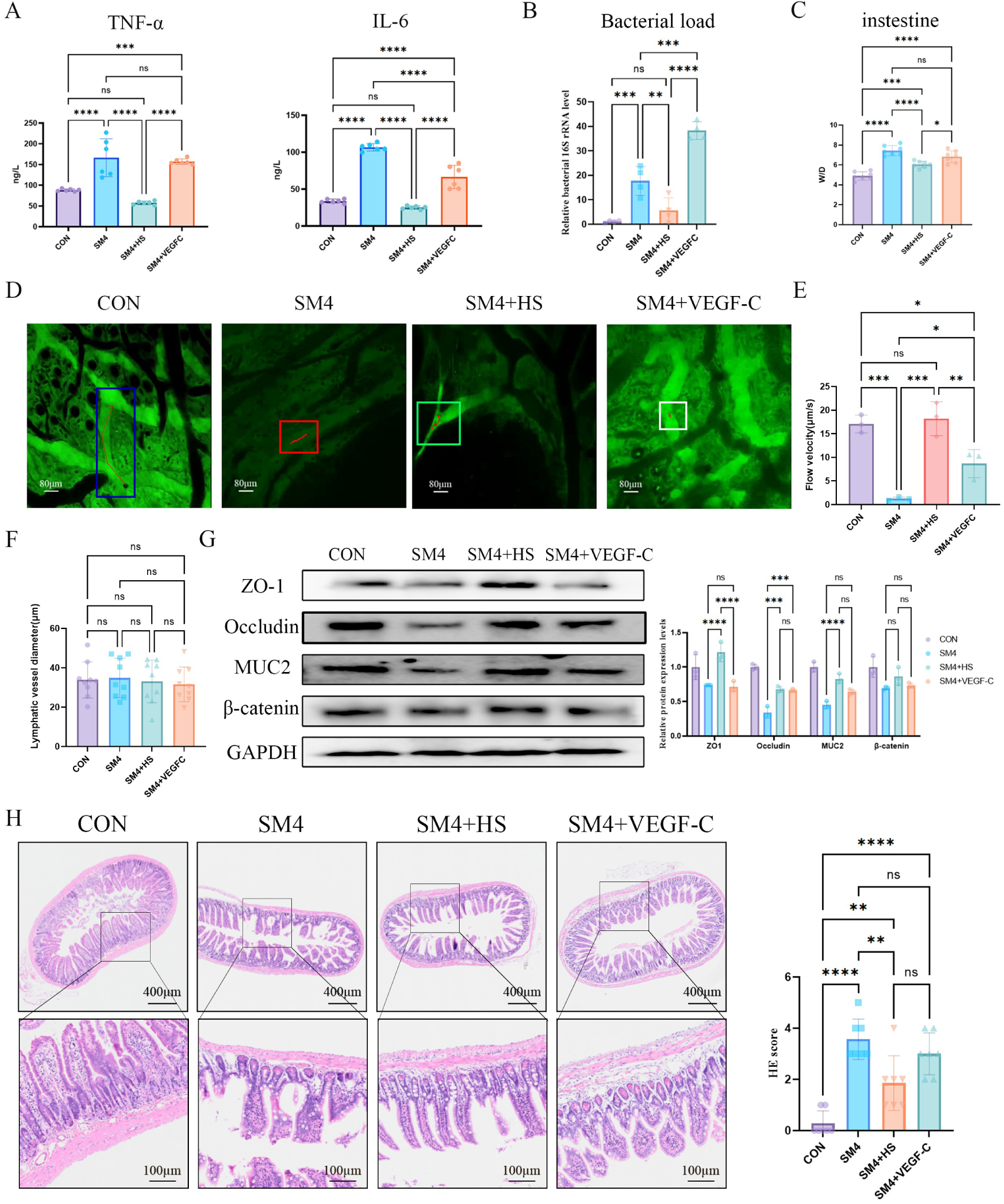
Inhibiting SOX18 transcription can trigger sepsis, while supplementing with HS has a protective effect. (A–B) SOX18 inhibitor SM4 elevates TNF-α/IL-6(n=6) and bacterial load(n=4). **(C)** Elevated W/D ratio indicating aggravated edema(n=6).**(D–E)** In vivo lymphatic imaging showing SM4-induced lymphatic reflux impairment, partially reversed by HS or VEGF-C supplementation. The curve marked in the box represents the movement trajectory of cells in the lymphatic vessels per unit time. **(F)** The diameter of the lymphatic vessels. **(G)** Relative expression of intestinal barrier proteins(n=3). **(H)** H&E staining and injury scoring of intestinal. Data are presented as mean ± SEM. *P < 0.05, **P < 0.01, ***P < 0.001, ****P < 0.0001; ns, not significant.

At the tissue level, SM4 treatment severely compromised intestinal barrier function, manifested by pronounced tissue edema and increased intestinal permeability(Figure 6C). This was accompanied by a significant downregulation of multiple key barrier-associated proteins, including ZO-1, Occludin, MUC2, and β-catenin(Figure 6G). In vivo functional imaging further revealed a substantial reduction in lymphatic flow velocity following SM4 treatment, indicating impaired lymphatic reflux(Figure 6D–E; Video 3). Despite these functional deficits, no significant differences in lymphatic vessel diameter were observed among the experimental groups, suggesting that overall lymphatic vessel architecture remained morphologically intact(Figure 6F). These findings indicate that SOX18 primarily regulates lymphatic function rather than vessel morphology.

Consistently, histopathological analysis further confirmed marked disruption of intestinal tissue structure in SM4-treated mice(Figure 6H). Notably, supplementation with HS significantly ameliorated multiple SM4-induced abnormalities, including inflammatory cytokine production, intestinal bacterial burden, intestinal permeability, lymphatic flow, and barrier protein expression. In contrast, VEGF-C administration conferred partial protection, but its effects were consistently less pronounced than those achieved by HS supplementation. Collectively, these data demonstrate that disruption of the VEGFR3 – SOX18 – HSPG2 signaling pathway compromises lymphatic reflux and barrier integrity, leading to heightened inflammatory responses. These findings further establish the functional importance of this axis in preserving lymphatic homeostasis and preventing tissue injury.

### 7. Genetic variants of HSPG2 are strongly associated with lymphatic system disorders and sepsis risk in human populations

To further validate the impact of HSPG2 on the risk of lymphatic dysfunction and sepsis, we analyzed genetic variation within the HSPG2 locus in a large population-based cohort. A total of 73 single-nucleotide polymorphisms(SNPs) within the HSPG2 gene region were identified and systematically evaluated for disease association. Among these, the top 30 variants exhibiting the strongest associations were selected for downstream analyses. In the cohort of individuals with lymphatic system disorders(Figure 7A–B), HSPG2 variants showed strong associations with lymphatic pathology(Figure 7C). Several SNPs demonstrated markedly elevated risk(e.g., rs369899077, rs371225050), with odds ratios exceeding 2(Figure 7D), suggesting that structural or regulatory variants in HSPG2 may directly impair lymphatic vessel stability and transport function.

**Figure 7.**
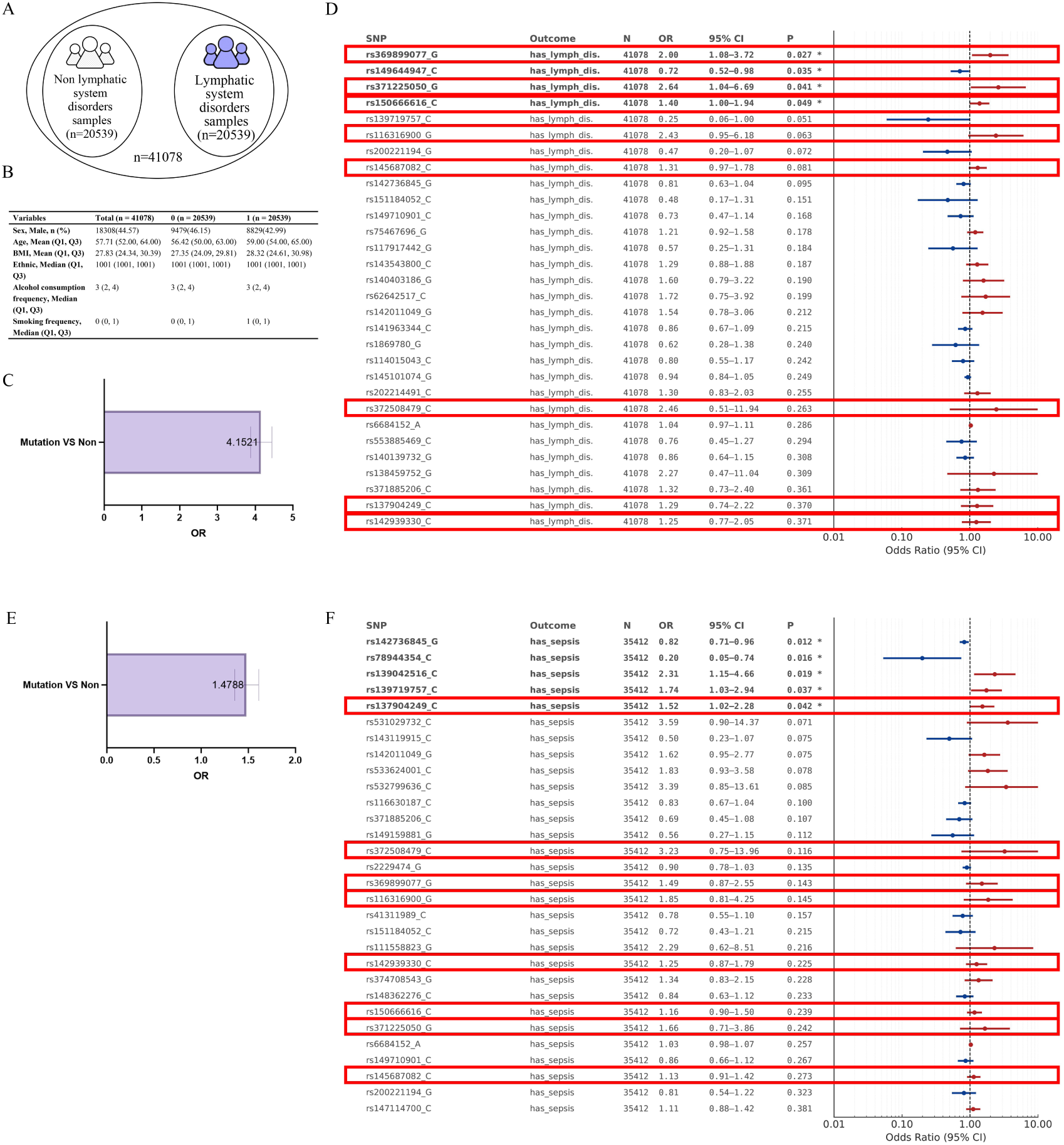
HSPG2 genetic variants are strongly associated with lymphatic disorders and sepsis. **(A)** Schematic diagram of lymphatic system disorder sample(n = 41,078). **(B)** Subject information. **(C)** Individuals carrying HSPG2 variants show a further increased risk of lymphatic system disorders(OR = 4.1521). **(D)** Top 30 HSPG2 SNPs ranked by association significance with lymphatic system disorders in the UK Biobank cohort. **(E)** Individuals carrying HSPG2 variants also exhibit an increased risk of sepsis(OR = 1.4788). **(F)** Top 30 HSPG2 SNPs ranked by association significance with sepsis in the UK Biobank cohort. Red boxes denote SNPs acting as shared risk variants in both the lymphatic disorder and sepsis cohorts. Data are presented as odds ratios(OR) with 95% confidence intervals(CIs).

We next examined the same SNP set in the sepsis cohort(n = 35,412). Interestingly, many HSPG2 variants also exhibited significant associations with sepsis(Figure 7E–F). Variants such as 139042516, 139719757 and 137904249 significantly increased the risk of sepsis(OR > 1.4), indicating that genetic perturbations of HSPG2 influence not only lymphatic homeostasis but also systemic host defense during infection. Strikingly, among the top 30 SNPs for each disease, 14 were shared between the two cohorts, and 12 of these exhibited concordant effect directions, with several SNPs acting as shared risk alleles(red boxes). These overlapping patterns strongly support a genetically conserved axis linking HSPG2 function to both lymphatic physiology and severe infection. Collectively, these findings demonstrate that genetic variation in HSPG2 is strongly associated with both lymphatic pathology and the occurrence of sepsis.

## Discussion

This study identifies lymphatic dysfunction as an active and previously underappreciated driver of sepsis pathogenesis. By integrating population-level genetic analyses, single-cell transcriptomics, and functional in vivo models, we demonstrate that impairment of lymphatic reflux is not merely a secondary consequence of systemic inflammation but represents a primary pathogenic event that actively promotes sepsis. Central to this process is the VEGFR3 – SOX18 – HSPG2 signaling axis, which governs lymphatic endothelial integrity and inflammatory homeostasis. Disruption of this axis compromises lymphatic structure and function, thereby predisposing tissues to uncontrolled inflammatory dissemination (Figure 8).

**Figure 8.**
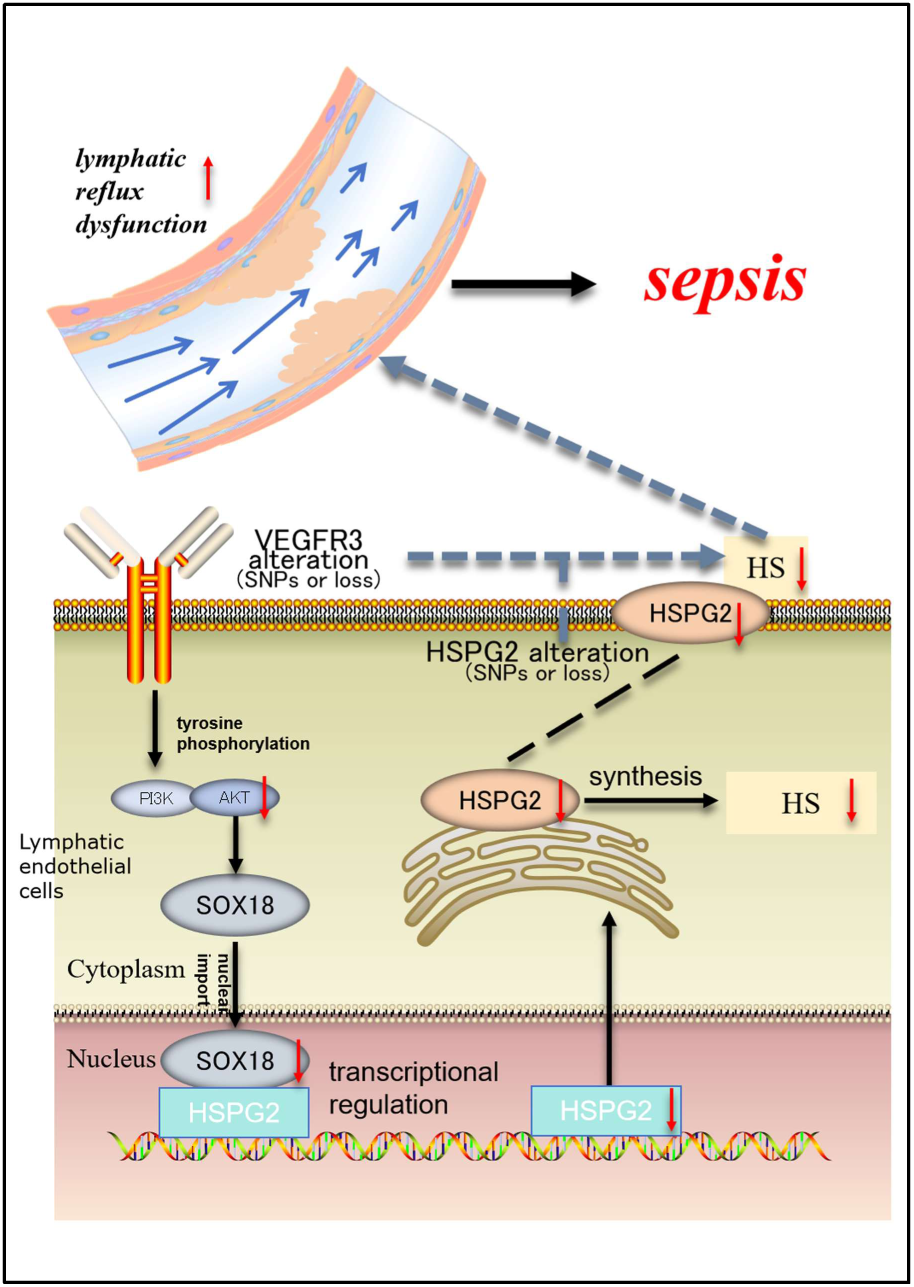
Schematic illustration of VEGFR3–SOX18–HSPG2 axis in regulating lymphatic reflux and sepsis. Genetic variants or functional loss of VEGFR3 impair SOX18 nuclear localization and reduce HSPG2-mediated HS synthesis in lymphatic endothelial cells, disrupting lymphatic reflux and barrier structures of lymphatic vessels, inducing sepsis.

Previous studies have shown that structural disruption of lymphatic vessels can provoke systemic inflammatory responses, and that fluid accumulation resulting from LECs injury is closely associated with adverse outcomes in sepsis^[4, 25]^. In parallel, enhancement of meningeal lymphatic drainage has been reported to alleviate disease severity in experimental models^[26]^. However, whether lymphatic dysfunction itself actively contributes to the initiation and progression of sepsis has remained unresolved. Here, we demonstrate that disruption of the VEGFR3–SOX18–HSPG2 signaling axis is sufficient to induce spontaneous sepsis, whereas restoration of this pathway—particularly through exogenous supplementation of heparan sulfate—markedly ameliorates disease manifestations. Together, these findings support a causal role for lymphatic dysfunction in sepsis.

SOX18 has been widely recognized as a central regulator of lymphatic endothelial identity and stability, acting through direct transcriptional control of genes involved in extracellular matrix organization and endothelial barrier maintenance, its related gene features are considered prognostic markers for sepsis^[27–29]^. Previous studies have shown that SOX18 binds promoter and enhancer regions of lymphatic-specific genes to coordinate structural and functional programs within lymphatic endothelium^[30, 31]^. In this context, our findings extend the known regulatory scope of SOX18 by linking its transcriptional activity to the expression of HSPG2. Through this transcriptional hierarchy, VEGFR3 signaling is positioned upstream of SOX18 to modulate the molecular composition of glycocalyx. This framework provides a mechanistic basis for how VEGFR3-dependent transcriptional regulation can influence lymphatic endothelial properties via HS-related components, setting the stage for subsequent alterations in lymphatic reflux function.

The lymphatic glycocalyx, enriched in heparan sulfate proteoglycans, constitutes a critical structural and functional interface that supports efficient lymphatic transport by reducing hydraulic resistance, preserving endothelial junctional integrity, and limiting nonspecific solute permeability^[21]^. Disruption of HS impaired, and reduced clearance of interstitial solutes^[32–34]^. In parallel, weakens the barrier properties of lymphatic endothelium, facilitating the translocation of microbial products and inflammatory mediators from peripheral tissues—particularly the intestine—into the systemic circulation, induce sepsis^[20]^.

Our findings provide convergent clinical and genetic evidence linking lymphatic dysfunction to sepsis. In population-level analyses of more than 500,000 individuals from the UK Biobank, variants in VEGFR3 were significantly enriched among individuals with sepsis, extending its relevance from lymphatic biology to clinical sepsis risk. Beyond VEGFR3, we further identified strong associations between HSPG2 genetic variants and both lymphatic system disorders and sepsis, suggesting that alterations in a key glycocalyx scaffold may represent a shared molecular basis underlying these phenotypes. Together, these data support a model in which inherited disruption of lymphatic regulatory pathways, first reflected by VEGFR3 variation and further captured by HSPG2 locus signals, is linked to clinically observable sepsis. Mechanistically, these genetic associations align with our experimental evidence that perturbation of the VEGFR3–HSPG2 axis compromises lymphatic transport capacity, thereby facilitating pathological dissemination of inflammatory mediators and contributing to sepsis.

In conclusion, this study identifies a VEGFR3–SOX18–HSPG2 signaling axis as a critical regulator of lymphatic reflux and inflammatory control. By linking lymphatic endothelial dysfunction to sepsis pathogenesis, our work highlights lymphatic vessels as active participants in sepsis and suggests new avenues for therapeutic intervention targeting lymphatic biology.

## Materials and Methods

### 1. Clinical and genetic data analysis

#### 1.1 Data sources

Population-level data were obtained from the UK Biobank, comprising a total of 502,137 participants, all of whom were initially included in the analysis. Disease phenotypes were defined based on International Classification of Diseases, Tenth Revision (ICD-10) codes extracted from hospital episode statistics and related health records. Sepsis cases were identified using ICD-10 codes A40 and A41. Lymphatic system – related disorders were defined primarily using ICD-10 codes reflecting lymphatic dysfunction, including I89 (noninfective disorders of lymphatic vessels) and Q82.0 (hereditary lymphedema). Additional ICD-10 codes associated with lymphatic system disorders(C77, C46.3, D86.1, D86.2, I89, R59) were evaluated in exploratory analyses. Matched control individuals were selected from participants without corresponding ICD-10 diagnoses. Genotype data were derived from the UK Biobank genotyping array dataset. Single-nucleotide polymorphism (SNP) annotation and reference rsID information were obtained from the Ensembl database(https://www.ensembl.org).

#### 1.2 Genetic statistical analysis

Statistical analyses were performed using R software(v4.3.2). Associations between SNPs and disease phenotypes were assessed using logistic regression models adjusted for age, sex, BMI, smoking status and alcohol consumption. Odds ratios(OR) and 95% confidence intervals were calculated using the glm() function. Results were summarized and visualized using forest plots.

### 2. In vivo experiments

#### 2.1 Animal models

C57BL/6 mice were used for all in vivo studies. Animals were housed under specific pathogen-free(SPF) conditions at 22 ± 2 °C with 50% ± 5% humidity, provided food and water ad libitum, and maintained on a 12-h light/dark cycle. All animal procedures were approved by the Animal Ethics and Welfare Committee of Wenzhou Medical University(Approval No. Wydw 2021-0182).

##### 2.1.1 In Vivo Pharmacological Treatments

For pharmacological intervention experiments, mice were treated with a SOX18 inhibitor(SM4) (20 mg/kg, intraperitoneal injection)^[35]^. SM4 was purchased from GLPBIO. To evaluate rescue effects, HS(50 mg/kg, intraperitoneal injection; GLPBIO)^[36]^ or recombinant VEGF-C (Cys156Ser) protein (0.1 μg/g body weight, intraperitoneal injection; R&D Systems)^[37]^ was administered 6 h after SM4 treatment. All reagents were freshly prepared and delivered in a total injection volume of 100–200 μL per mouse.

##### 2.1.2 Generation of conditional knockout mice

1. Prox1-CreERT2 mice were crossed with VEGFR3^flox/flox(Shanghai Model Organisms Center) mice to generate VEGFR3^flox/flox;Prox1-Cre mice(hereafter abbreviated as VEGFR3^iΔLEC). Mice were randomly assigned to groups and administered tamoxifen(75 mg/kg, i.p.) for five consecutive days to induce gene deletion. Control mice carried the same genotype (VEGFR3^flox/flox;Prox1-Cre) but did not receive tamoxifen, thereby serving as tamoxifen-untreated controls.
2. AAV-CMV>DIO-mHSPG2-Kozak vectors were custom-generated and packaged into AAV9 serotype(Cyagen, China). Following tamoxifen induction, Prox1-Cre mice received an i.p. injection of the virus(2×10^11 GC/mouse), as shown in Fig.4A. Control mice received an equal titer of AAV-scramble. Tissues were harvested 20 days after viral injection, based on the established time window required for AAV-mediated gene expression.

#### 2.2 Single-cell transcriptome analysis

Single-cell RNA-seq data were obtained from a CLP-induced sepsis mouse model(Sham vs. CLP; GEO accession GSE207651). Lung tissues were collected from sham-operated mice (0 h) and CLP-treated mice at 24 h post-surgery for sequencing. Data were processed using R(v4.3.2) for quality control, normalization, dimensionality reduction(t-SNE/UMAP), and clustering. LECs were annotated based on canonical markers, including Prox1, LYVE1, and Pdpn.

#### 2.3 Measurement of inflammatory cytokines

Whole blood was collected from each group, and serum TNF-α and IL-6 levels were measured using ELISA kits(Boyun Biotech, Shanghai) following the manufacturer’s instructions.

#### 2.4 Histological analysis

Tissues were fixed in 4% paraformaldehyde, paraffin-embedded, sectioned, and stained with hematoxylin and eosin(H&E). Intestinal villi morphology and lung structure were examined by light microscopy. Tissue edema was quantified by calculating the wet/dry weight ratio, a standard metric for assessing fluid accumulation.

#### 2.5 In vivo two-photon imaging of lymphatic flow

Lymphatic flow dynamics were visualized using two-photon intravital microscopy with an FVMPE-RS system (Olympus). After anesthesia with sodium pentobarbital(1%), mice were depilated and their intestines gently exposed. FITC-conjugated dextran(2000 kDa; Invitrogen) was injected into Peyer’s patches to label lymphatic vessels. To minimize motion artifacts caused by respiration and muscle contraction, the intestine was elevated using an adsorption-based imaging holder, allowing the tissue to be positioned away from surrounding muscle structures. Time-lapse imaging was performed to record lymphatic flow. Flow velocity and cellular trajectories were quantified using Python (OpenCV) based on manual particle tracking. Lymphatic vessel diameter was measured using Image J.

#### 2.6 Immunofluorescence

Frozen tissue sections were prepared and equilibrated to room temperature. After blocking with 5% donkey serum, sections were incubated overnight at 4 °C with primary antibodies(rabbit). Following washing, Alexa Fluor-conjugated donkey anti-rabbit secondary antibodies were applied for 1 h at room temperature(protected from light). Nuclei were counterstained with DAPI and samples imaged using confocal microscopy.

### 3. In vitro experiments

#### 3.1 Cell culture and transfection

##### 3.1.1 siRNA knockdown and plasmid construction

Four siRNAs targeting VEGFR3(IDs 3302, 3307, 3318, 3387; sequences in Supplementary Table 1) were synthesized by GenePharma. VEGFR3 and SOX18 overexpression plasmids were constructed. HSPG2 promoter fragments(−200, −441, −500, −1000 bp) were synthesized by Neupro Biotech for subsequent promoter activity assays(sequences in Supplementary Table 2).

##### 3.1.2 HLEC culture and model establishment

Human lymphatic endothelial cells(HLECs; Chinese Academy of Sciences Cell Bank) were cultured in 1001 Endothelial Cell Medium(ScienCell, USA). Transfection was performed using Lipofectamine 3000(Thermo Fisher Scientific, USA).

#### 3.2 RNA extraction and quantitative real-time PCR

Total RNA was extracted using RNA-isolation reagent(Takara, Japan). cDNA synthesis was performed using HiScript® III All-in-One RT SuperMix(Vazyme, China). qPCR reactions were run using FastStart Universal SYBR Green Master(TORO, China) on a QuantStudio™ 3 system(Thermo Fisher, USA). GAPDH served as the reference gene, and relative mRNA expression was calculated via the ΔΔCt method. Primer sequences are listed in Supplementary Table 3.

#### 3.3 Western blotting

Proteins were extracted using RIPA buffer with PMSF(Beyotime, China). Samples were separated by SDS-PAGE, transferred onto PVDF membranes, and blocked with 5% skim milk. Primary antibodies were applied overnight at 4 °C, followed by secondary antibodies(Beyotime) the next day. Detection was performed using ECL, and band intensities were quantified using Image J. Antibodies used included:

1. Rabbit IgG(#30000-0-AP), mouse IgG(#B900620), Proteintech
2. Anti-VEGFR3(#sc-514302), anti-SOX18(#sc-7383), anti-HSPG2(#sc-7345), Santa Cruz
3. anti-Occludin(DF7504), anti-beta catenin(AF6266), anti-MUC2(DF8390), anti-LYVE1(AF4202), anti-ZO1(AF5145), Affinity
4. Anti-GAPDH(#MB001), Bioworld

#### 3.4 High-throughput sequencing

Total RNA was isolated using Trizol, and sequencing libraries were constructed and sequenced on an Illumina platform. Differentially expressed genes were identified using DESeq2 with thresholds |log₂FC| > 0.58 and P < 0.05. Enrichment and functional annotation analyses were subsequently performed.

#### 3.5 Dual-luciferase reporter assay

Three HSPG2 promoter fragments(−1000, −500, −441 bp) were cloned into luciferase reporter vectors. Recombinant plasmids and Renilla controls were co-transfected into LECs. After 48 h, luciferase activities were measured sequentially, and relative transcriptional activity was calculated(Promega, China).

#### 3.6 Chromatin immunoprecipitation(ChIP)

Cells were cross-linked with 1% formaldehyde. Nuclei were isolated, and chromatin was sheared to 200–600 bp fragments using an automated sonicator. Samples were divided into IP(0.8 mL), IgG control(0.8 mL), and Input(0.1 mL) groups. IP samples were incubated with 3–8 µg of SOX18 antibody, and IgG controls received an equal amount of IgG. After overnight rotation at 4 °C, Protein A/G magnetic beads were added for 30 min. Beads were washed, eluted, and subjected to reverse cross-linking at 65 °C for 6 h. Purified DNA was analyzed by ChIP-qPCR(BersinBio, China).

### 4. Statistical analysis

All experimental results were repeated at least three times. Data are presented as mean ± SEM. Two-group comparisons were performed using two-tailed Student’s t-tests, while multiple-group comparisons were analyzed using one-way ANOVA. Survival analyses were performed using Kaplan–Meier curves and log-rank tests. P < 0.05 was considered statistically significant. Statistical analyses were conducted using GraphPad Prism 9.0(San Diego, CA, USA) and R software.

## Acknowledgments

We sincerely acknowledge the support from the Oujiang Laboratory(Wenzhou, China). We also thank the Research and Experimental Center of Wenzhou Medical University for providing technical assistance and instrument access during this study.

---

**Supplementary Table 1.**
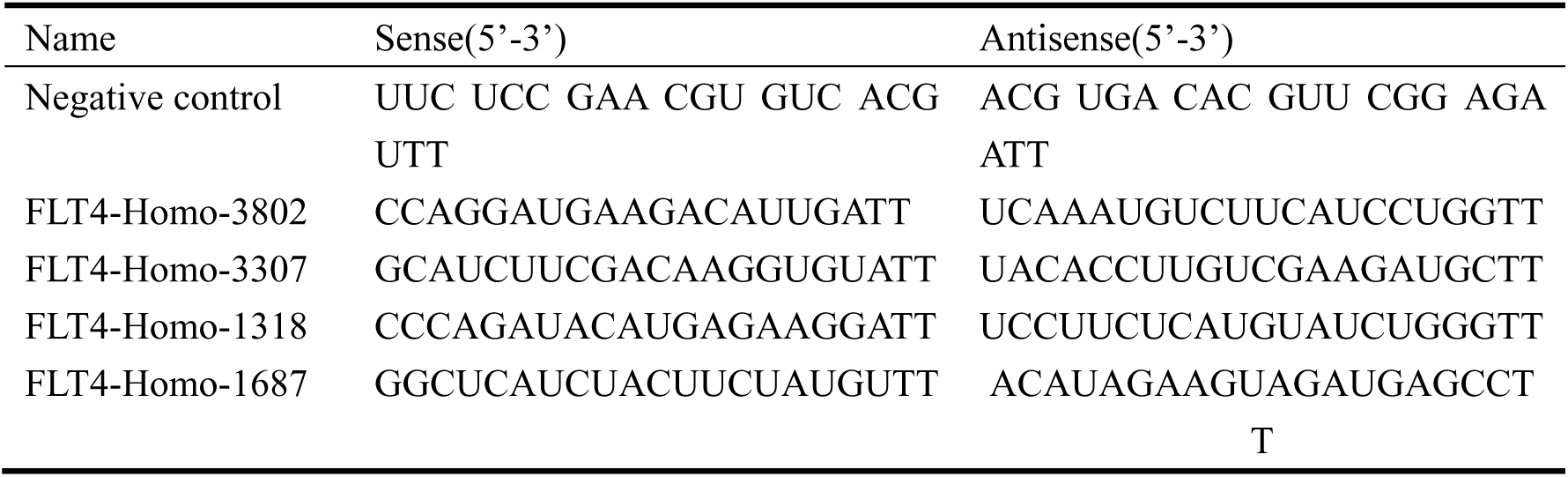
Sequence of siRNA.

**Supplementary Table 2.**
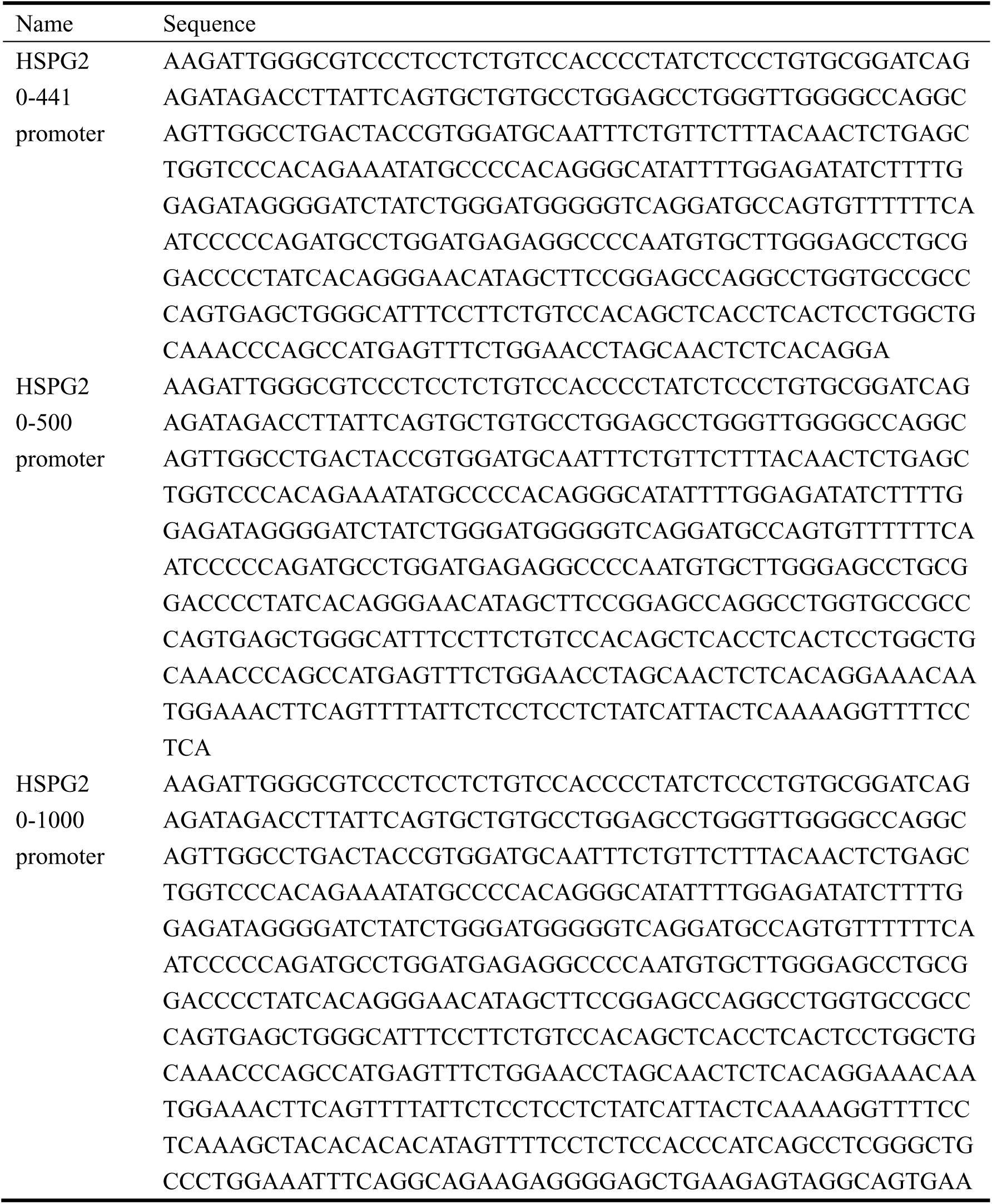
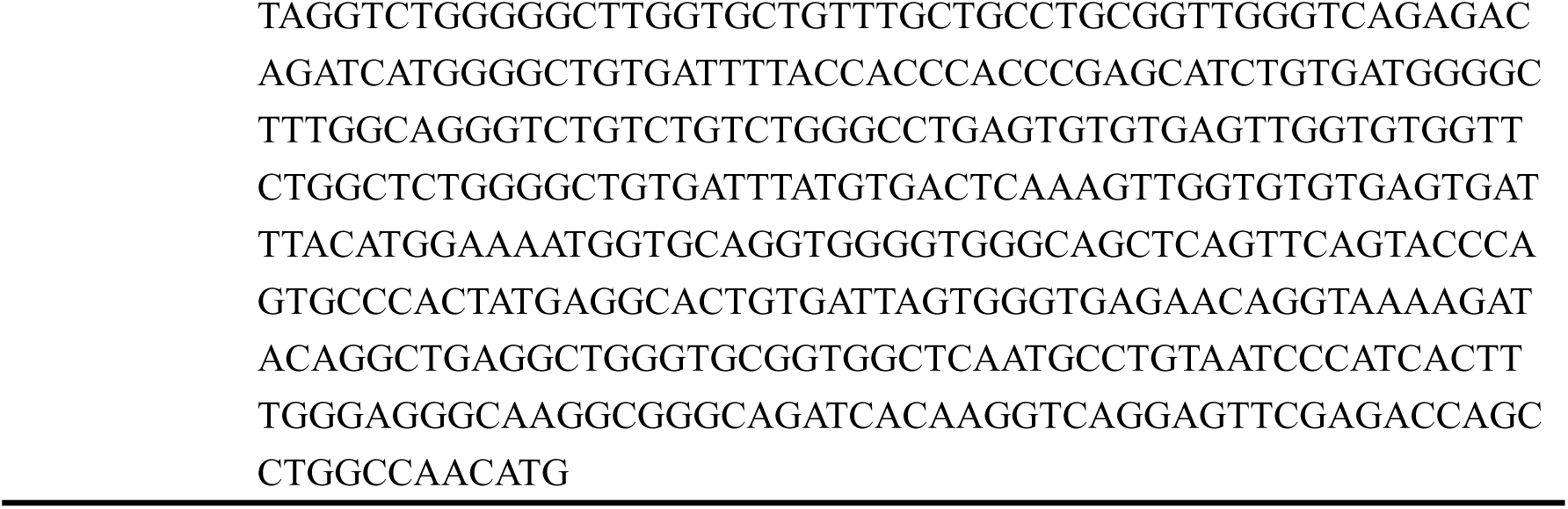
Sequence of HSPG2 promoter fragments.

**Supplementary Table 3.**
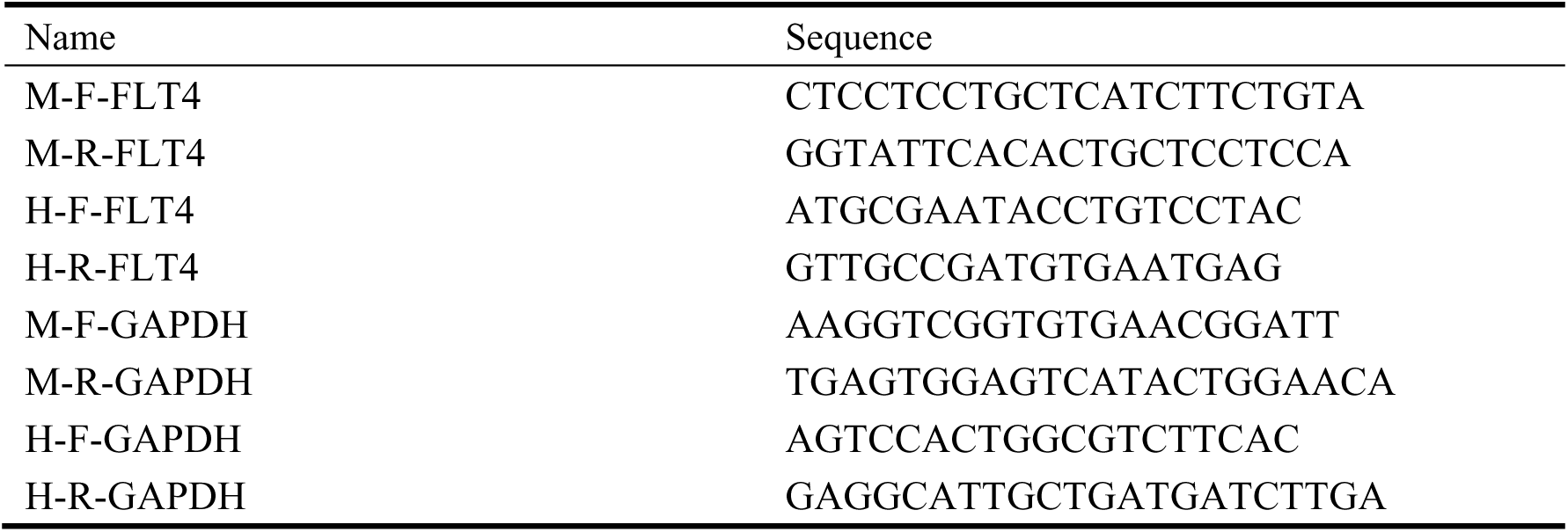
Primer sequence.

